# Analysis and Forecasting of Global RT-PCR Primers for SARS-CoV-2

**DOI:** 10.1101/2020.12.26.424429

**Authors:** Gowri Nayar, Edward E. Seabolt, Mark Kunitomi, Akshay Agarwal, Kristen L. Beck, Vandana Mukherjee, James H. Kaufman

## Abstract

Rapid tests for active SARS-CoV-2 infections rely on reverse transcription polymerase chain reaction (RT-PCR). RT-PCR uses reverse transcription of RNA into complementary DNA (cDNA) and amplification of specific DNA (primer and probe) targets using polymerase chain reaction (PCR). The technology makes rapid and specific identification of the virus possible based on sequence homology of nucleic acid sequence and is much faster than tissue culture or animal cell models. However the technique can lose sensitivity over time as the virus evolves and the target sequences diverge from the selective primer sequences. Different primer sequences have been adopted in different geographic regions. As we rely on these existing RT-PCR primers to track and manage the spread of the Coronavirus, it is imperative to understand how SARS-CoV-2 mutations, over time and geographically, diverge from existing primers used today. In this study, we analyze the performance of the SARS-CoV-2 primers in use today by measuring the number of mismatches between primer sequence and genome targets over time and spatially. We find that there is a growing number of mismatches, an increase by 2% per month, as well as a high specificity of virus based on geographic location.

## Introduction

As the SARS-CoV-2 pandemic grows, an essential method for controlling its spread and determining readiness for the re-opening of public life is through rapid testing. Rapid tests for active SARS-CoV-2 infections are based on reverse transcription polymerase chain reaction (RT-PCR). These tests consist of a forward primer, reverse primer, and probe that together are used to amplify the signal from the targeted virus within a sample. The approach supports rapid and specific identification of the virus, and does not depend on tissue culture or animal cell models. However, RNA viruses evolve over time and a specific PCR test may lose sensitivity as the genotypic distribution of the virus changes or shifts. Phylodynamic studies suggest the mutation rate of SARS-CoV-2 is in the range 1.05 × 10^−3^ to 1.26 × 10^−3^ substitutions per site per year, approximately 1.5% variation increase per month,^1^ consistent with mutation rates reported for other *Coronaviridae*.^2–4^

Sequence drift also leads to geospatial differences in the virus, resulting in varying test sensitivity by region. This study investigates the effectivity of current SARS-CoV-2 PCR tests over the development of the virus in space and time, and projects how the performance of each may change as the virus undergoes mutation. By taking a global perspective, using specific PCR protocols from several different countries together with genomic data from around the globe, our analysis shows how the existing tests respond differently over both time and location. By analyzing the number of mismatches of the PCR primers with respect to the sequenced SARS-CoV-2 genomes, we can measure how the targeted proteins are mutating. This provides an understanding of possible shortcomings of current tests, and suggests how often we may need to update those tests in the future. Through this work, we observe an average rate of amino acid sequence change of approximately 3% per month for the targeted proteins. Furthermore, we see that the virus genotype is spatially differentiated to the point that inter-country PCR testing already leads to a much higher rate of mismatches.

In support for global pandemic response, several countries have published their RT-PCR protocols. We have collected the primer sequences and protocols developed for six different regions – USA, Germany, China, Hong Kong, Japan, and Thailand – as provided by the WHO^5^. For all six protocols, we collect the forward, reverse, and probe sequences for each specific gene target. Table 1 details the different gene targets for each protocol. Most commonly, the PCR tests target the nucleoprotein (NP), followed by targets in the RNA-directed RNA polymerase (RdRP) gene, and the envelope small membrane protein (E protein). NP is a structural protein that encapsidates the negative strand RNA. For other RNA viruses including influenza, the NP sequence is often used for species identification^6^. RNA-dependent RNA polymerase (RdRP) is an enzyme that catalyzes the replication of RNA from an RNA template. The membrane associated RdRP is an essential protein for *Coronavirus* replication^7^, and may be a primary target for the antiviral drug remdesivir^8^. The E protein is a small membrane protein involved in assembly, budding, envelope formation, and pathogenesis^9^. The SARS-CoV E protein also forms a Ca2+ permeable ion channel that alters homeostasis within cells which leads to the overproduction of IL-1beta^10, 11^.

**Table 1.**
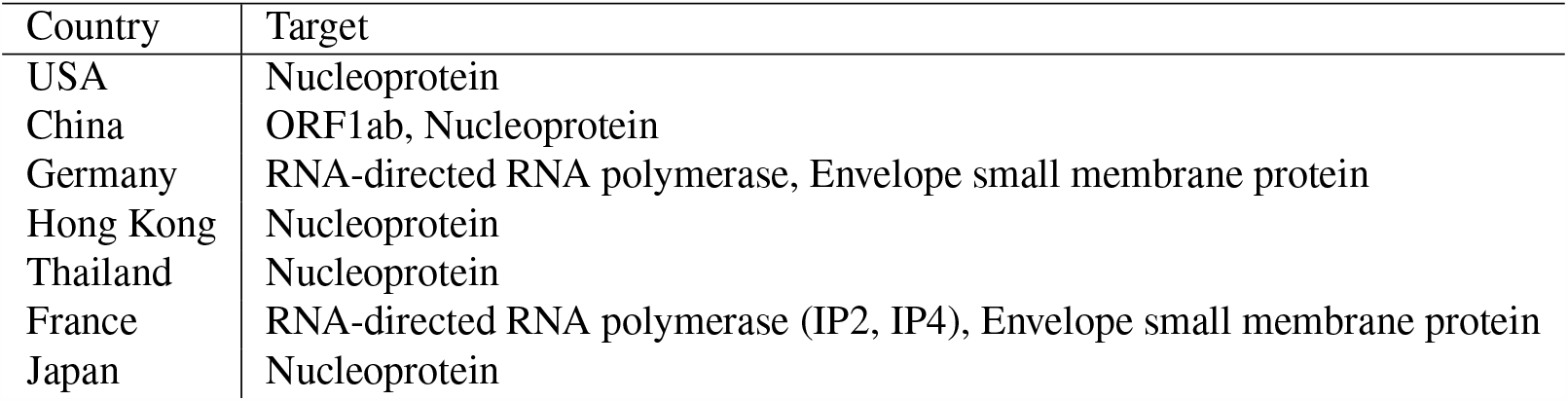
Targeted genes by name by primers from the countries in the study

| Country | Target |
| --- | --- |
| USA | Nucleoprotein |
| China | ORF1ab, Nucleoprotein |
| Germany | RNA-directed RNA polymerase, Envelope small membrane protein |
| Hong Kong | Nucleoprotein |
| Thailand | Nucleoprotein |
| France | RNA-directed RNA polymerase (IP2, IP4), Envelope small membrane protein |
| Japan | Nucleoprotein |

## Results

### 1 Primer Comparison

Using these methods, we observed high sequence homology for at least 95% of all genomes for most of the PCRs, showing that each primer is able to detect most of the SARS-CoV-2 genomes sequenced at the time of this report. Table 2 shows the percent of genomes hit by each PCR test, labelled by the country and target gene region. The America RP is an additional primer/probe set to detect the human RNase P gene to control for non-viral genes in the sample, and therefore, as expected, 0% of the SARS-CoV-2 genomes match with this set. However, when we look at the number of mismatches for each PCR for those hit genomes, we can see that there is a significant difference in performance between each test. Figure 1 shows the number of mismatches for all genomes created by each PCR, where we can see the range varying from 1796, created by the American N1 primer, to 42 mismatches, created by the French IP2 primer. Thus we observe that the measure of mismatches can be used as a proxy to identify the amount of variation found within the gene sequences that are being targeted by the worldwide tests.

**Table 2.**
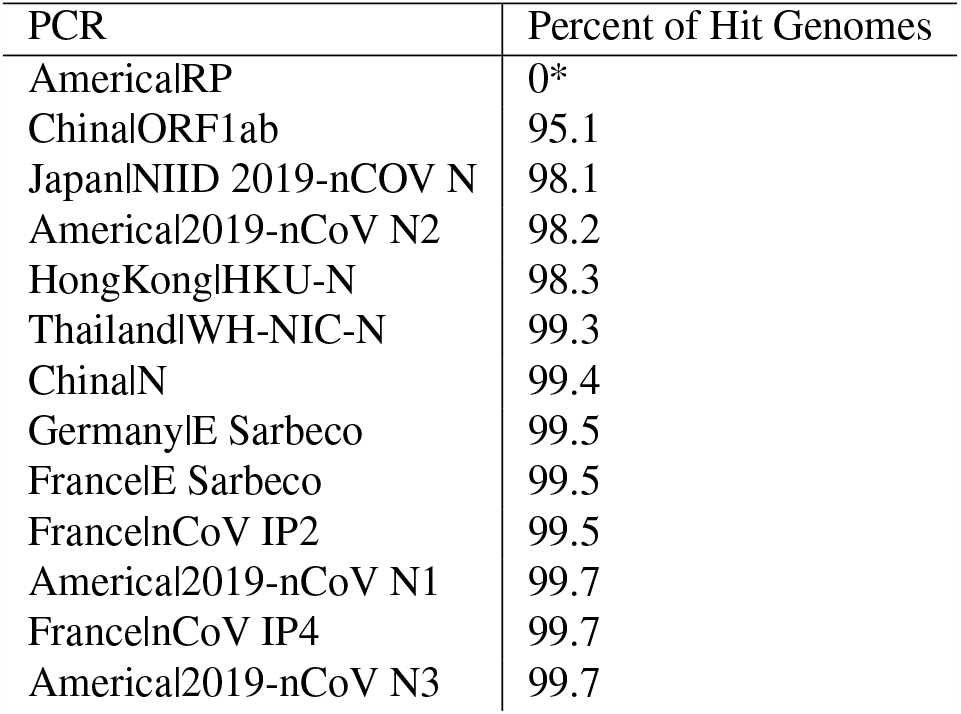
Percent of genomes that are hit by the described PCR test, identified by the country and target gene. *Indicates that the primer is designed to separate the any errant samples within the assay.

**Figure 1.**
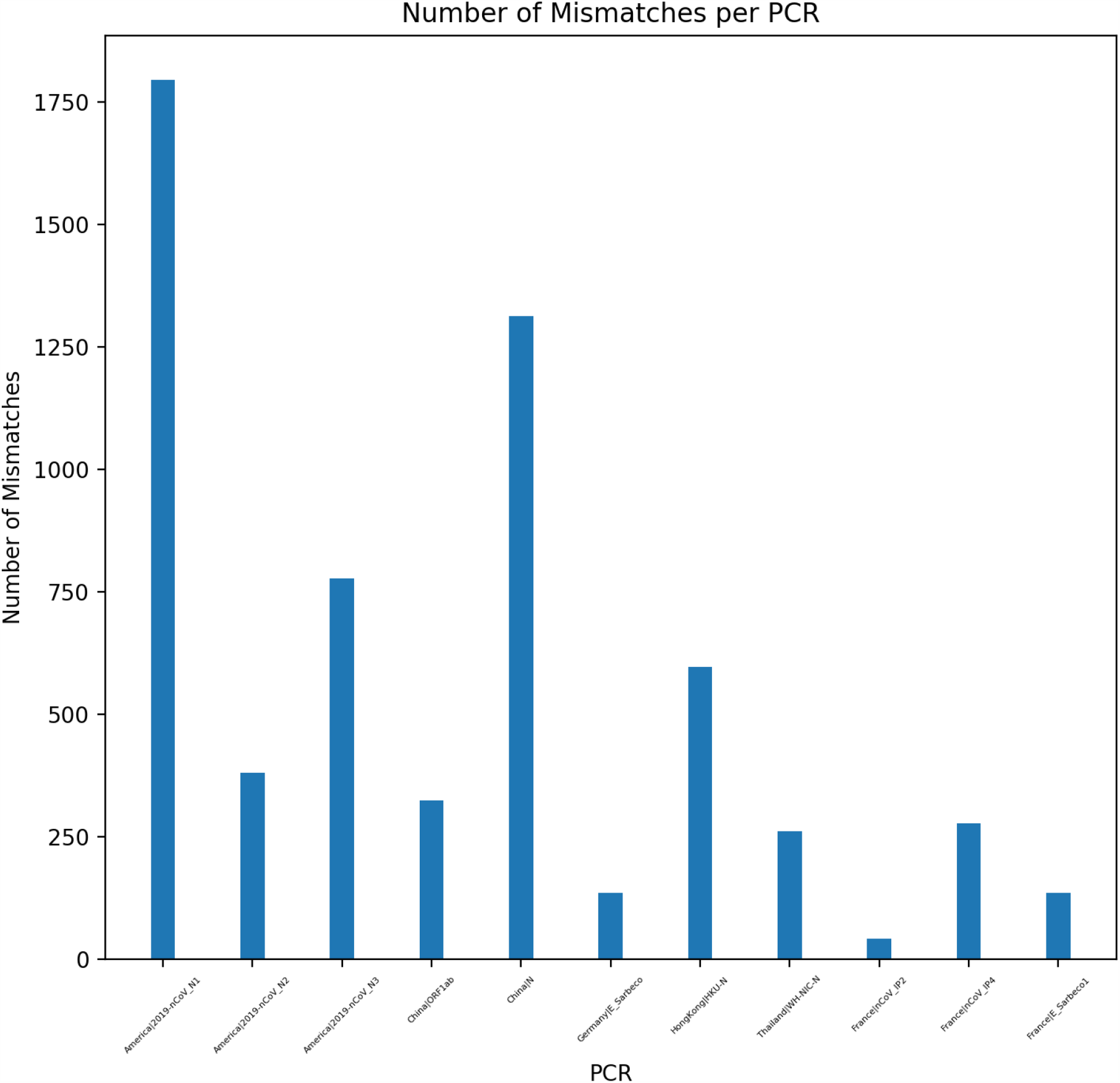
Total number of mismatches each PCR test creates when tested against the full corpus of SARS-CoV-2 genomes. Each PCR test is identified by the country of use and the targeted gene name.

### 2 Time Analysis

Following the methods described in Section 7, all genomes that fall within the 207 day range are segmented by date of collection and analyzed for mismatches to the various primer tests. Figure 2 shows the average number of mismatches seen for all primers each day within this range, normalized by the number of genomes sampled in each day. From this analysis, we can see an average of 1.1 mismatches, with a 14% increase in mismatches over the 207 day time range. This corresponds to a ~2% increase per month. To estimate the mutation rate,from figure 2, we calculate the best-fit line using least squares, which results in an R^2^ value of 0.6. This mutation rate is consistent with the expected rate of mutation of the SARS-CoV-2 virus.^1^ –4 Figure 3 shows the distribution of total, and time averaged, mismatches for each primer set over time. The figure indicates a larger distribution of mismatches for primer sets that target nucleoprotein regions.

**Figure 2.**
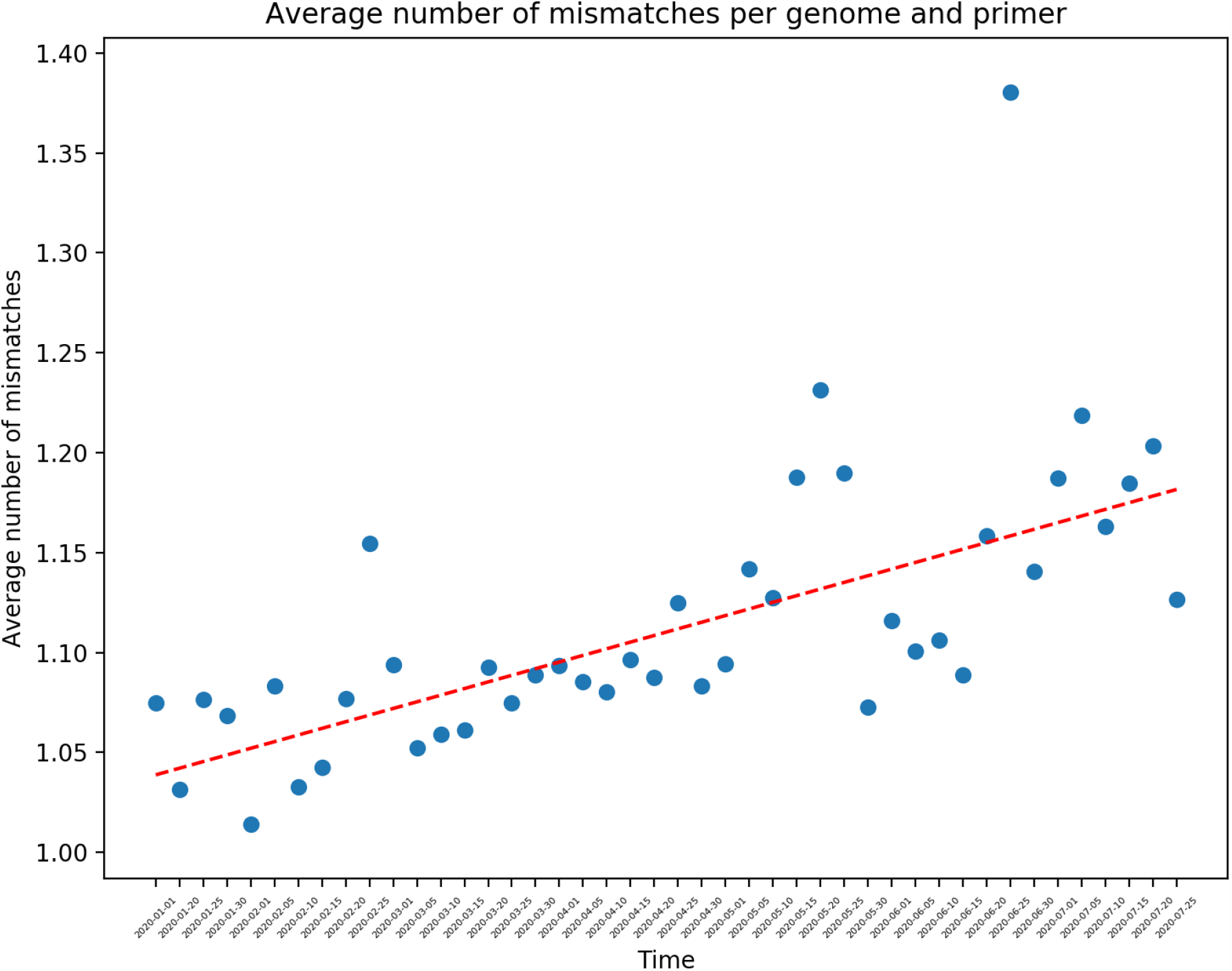
Average number of mismatches for all genomes and all PCR primers separated by the day on which the genome is collected. The dates shown are aggregated over every 5 day period.

**Figure 3.**
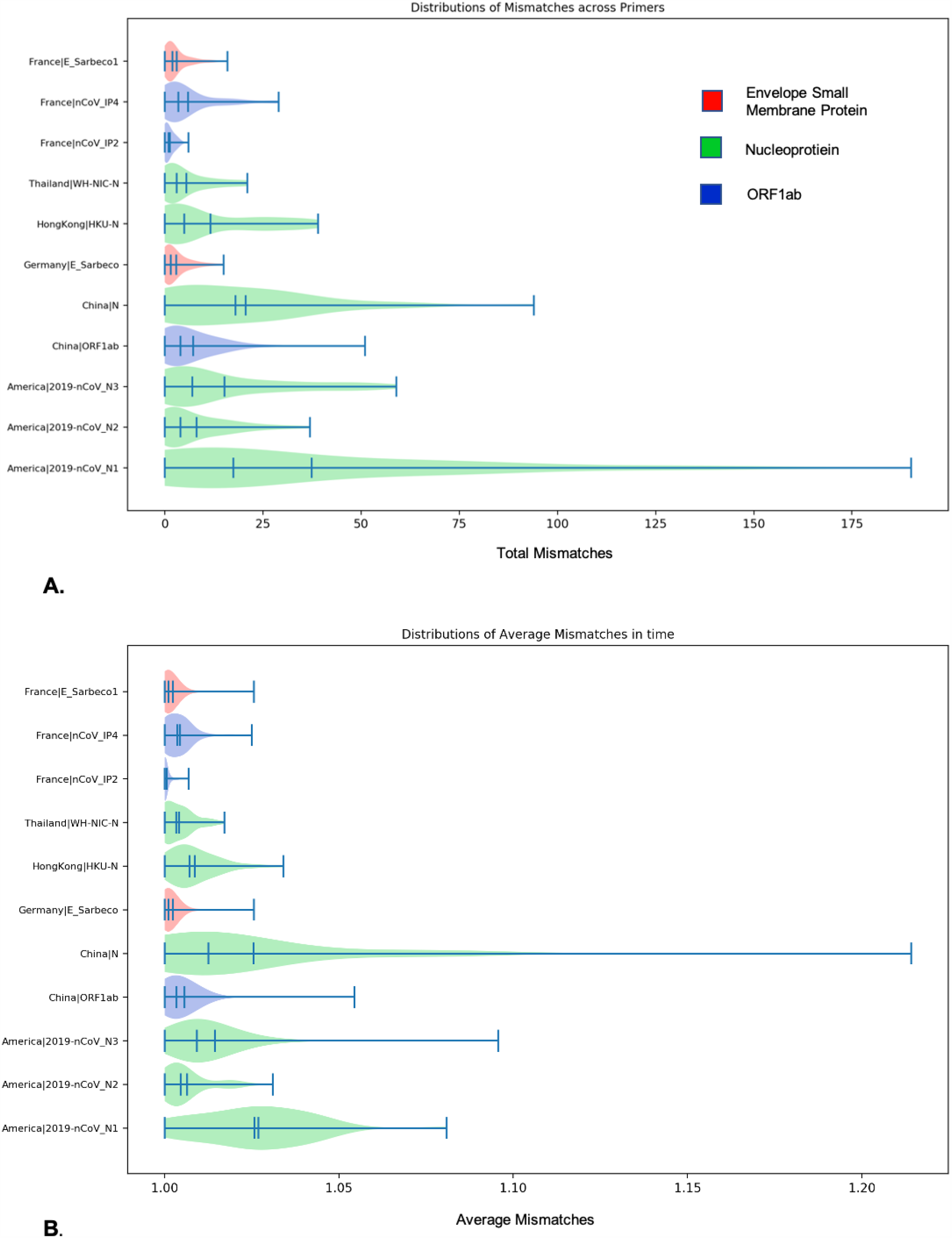
Distribution of mismatches for each primer. A shows the total number of mismatches aggregated for each day within the time range. B shows the number of mismatches for each day averaged by the number of genomes that occur on a day within the time range.

It is important to note that the total number of mismatches occurring is increasing and that many of these mismatches are being sustained in the evolving population. In order to identify a trend, genomes that occur close in time should have smaller change in mismatches than genomes that occur further apart in time. Figure 4 shows this comparison between delta time and delta mismatches for every pair of genomes for the France PCR targeting the RdRP gene (IP4). The graphs for the other PCRs may be found in the supplemental files. Each point represents a pairwise comparison of the difference in mismatch plotted over the difference in time. We observe that the delta mismatches grows in variance as the genomes occur further apart in time. Furthermore, the Pearson coefficient is 0.99 between mismatches and the number of genomes sampled in a day for each PCR. This positive linear relationship between the number of genomes and the number of mismatches per day shows that the mismatches occur uniformly across the genomes sampled within a day (rather than a few genomes creating noise in the signal). The data indicates that the virus demonstrated sequence variability in the targeted gene regions and that this variability causes sequence mismatches to increase over time.

**Figure 4.**
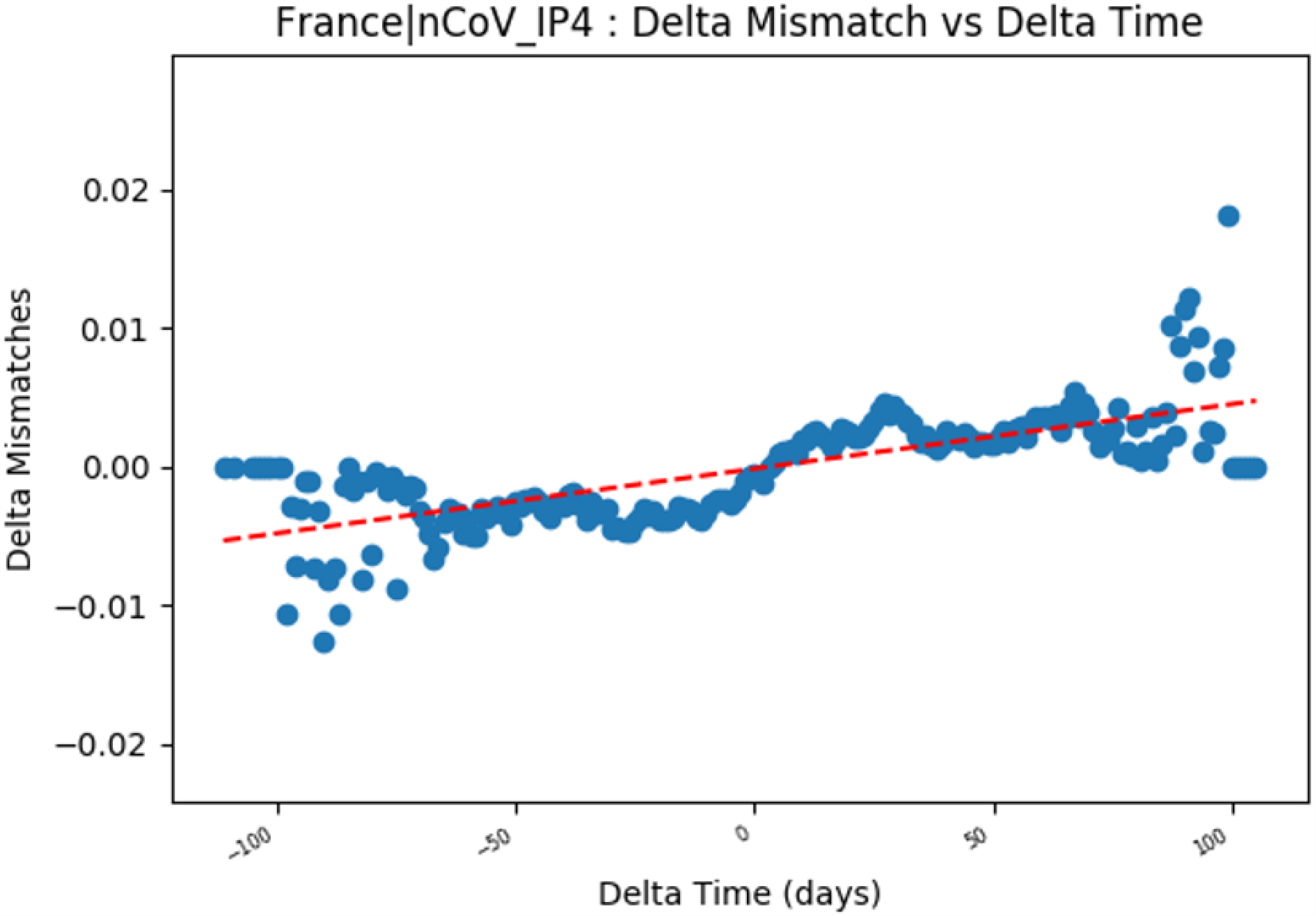
Change in number of mismatches between two occurrences over delta time between the two occurrences for the IP4 primer developed in France. The increasing slope shows that mutations are being sustained as we compare genomes that occur further apart in time. Graphs for all primers are included in the supplement.

### 3 Geographical Analysis

Geographical stratification is occurring as the SARS-CoV-2 virus mutates within each geographic location. Following the methods described in Section 8, geospatial analysis is conducted to identify patterns in mismatches found in genomes sequenced within versus outside the country of primer origin. Figure 5 shows the number of mismatches, normalized by the number of genomes within each category, for each PCR, grouped by same and other countries. There are 9 countries in which the number of mismatches in the country is lower than the number of mismatches that occur with genomes sampled outside of the country. This shows that the virus displays localized tendencies within the targeted gene regions, in addition to the spike glycoprotein region. The two outliers, the Hong Kong and France primers, show a higher percent of mismatches within the country rather than from different countries. Figure 6 shows the average number of mismatches over time, grouped by the genomes sampled within and outside the country, for one American primer. While the in-country average number of mismatches shows low variability, the out-country average number of mismatches show an increasing diversity in these targeted regions. The full set of graphs for each PCR tested are available in the supplement.

**Figure 5.**
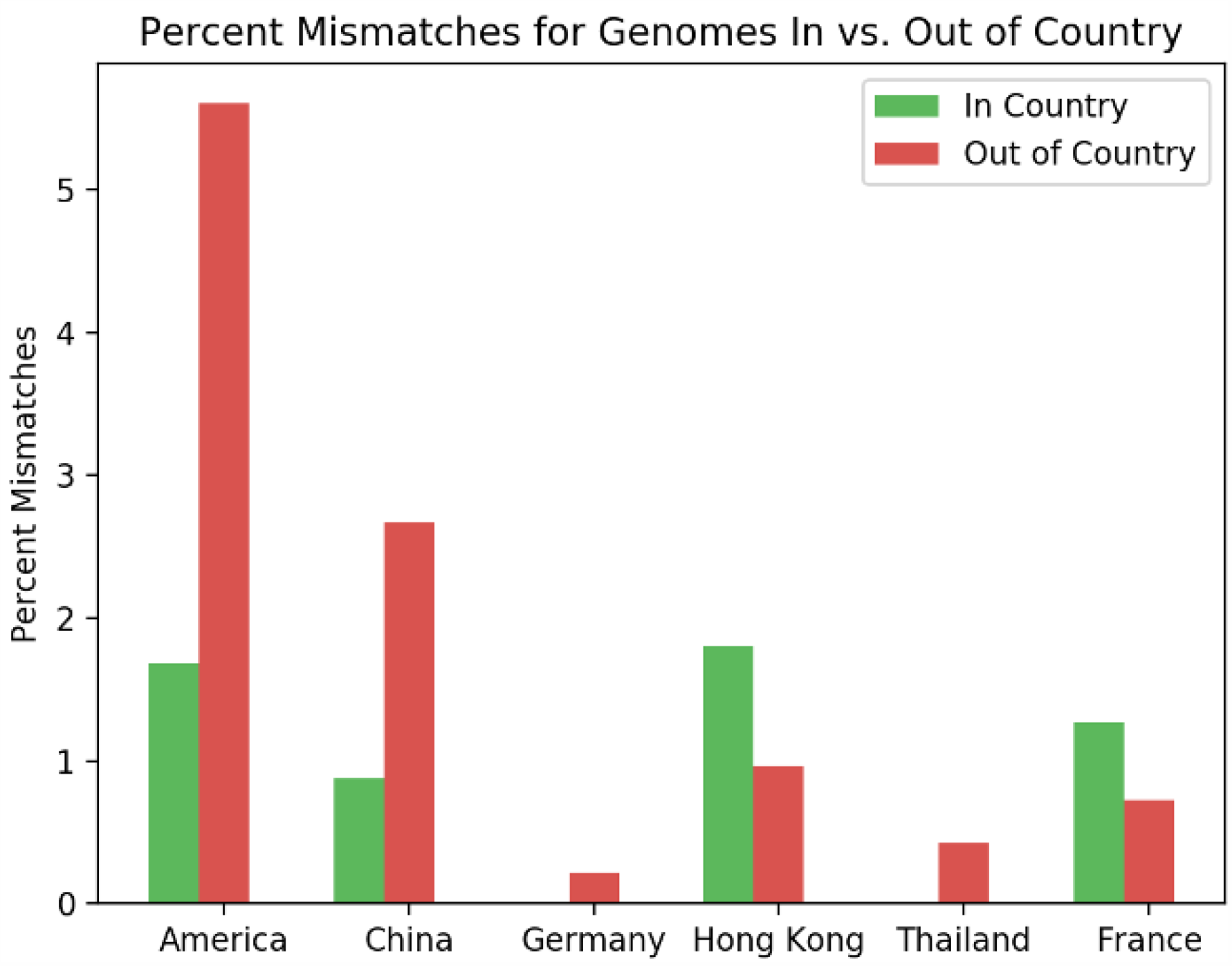
Number of mismatches for each PCR test tested on all SARS-CoV-2 genomes, split between genomes collected within the same country as the test and outside the country. For Japan, 100% of genomes, both in and out of the country, have 1 mismatch, and therefore not shown in the figure. For 9 out of the 11 PCR tests, there are a higher number of mismatches for total genomes that occur outside the country than genomes that occur inside the country.

**Figure 6.**
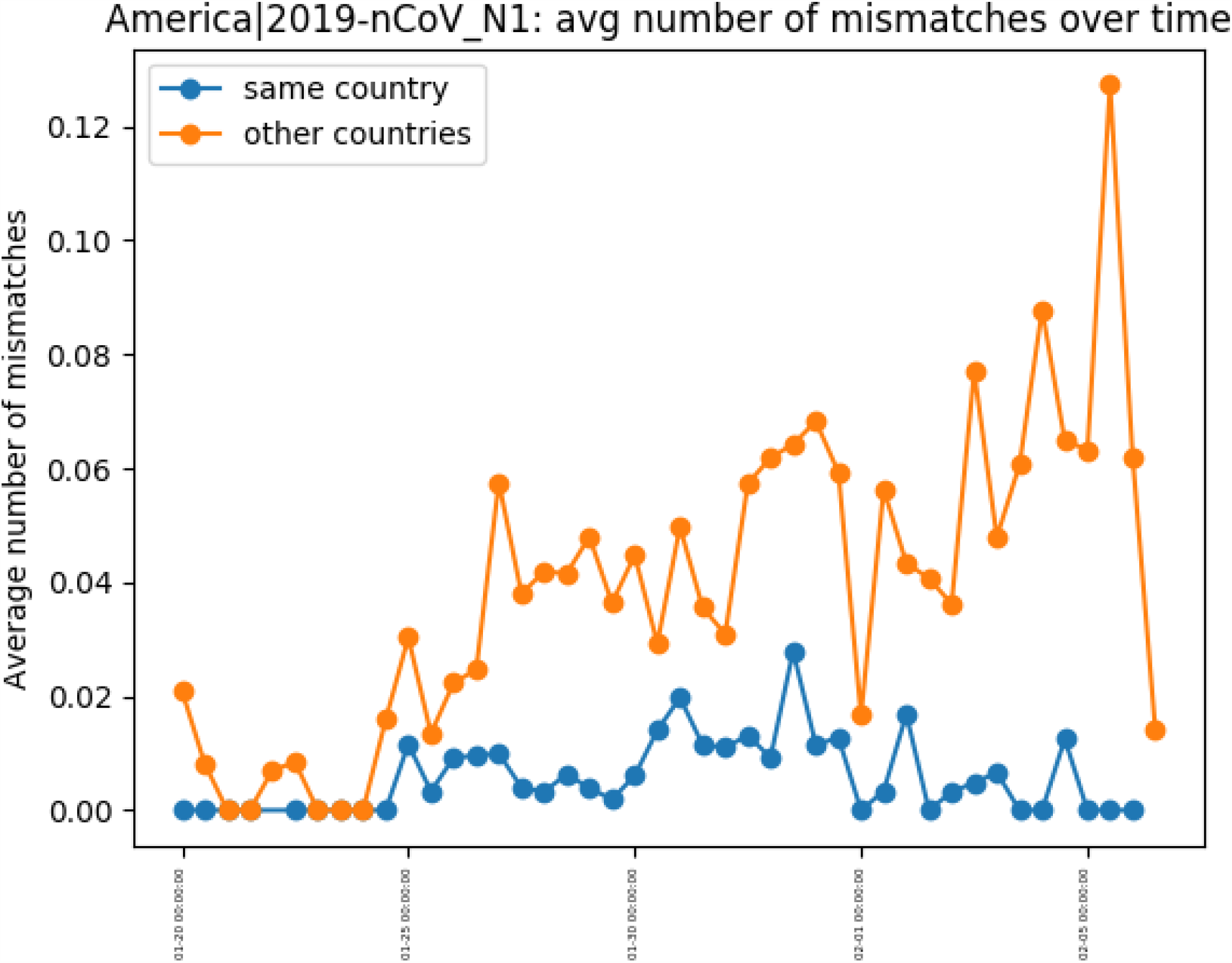
Number of mismatches in and out of country for an American nucleoprotein primer separated by time of genome collection. All other primers are included in the supplement.

### 4 Clade Analysis

Figure 7 shows the number of mismatches for each PCR per clade, normalized by the number of genomes in the PCR and clade. This shows definite trends which confirm the geographic specificity of the virus; for example, the American nucleoprotein primers have the highest number of mismatches for clade 19A, which Nextstrain defines as originating from predominantly Asian genomes, while the Chinese primer has the lowest number of mismatches for this clade. However, the clades are defined by specific mutations at nucleotide locations, which only overlaps with the primer bind region for 3.37% of the genomes. Therefore, the relationship between the primer mismatches and the genome clades are correlational rather than causational.

**Figure 7.**
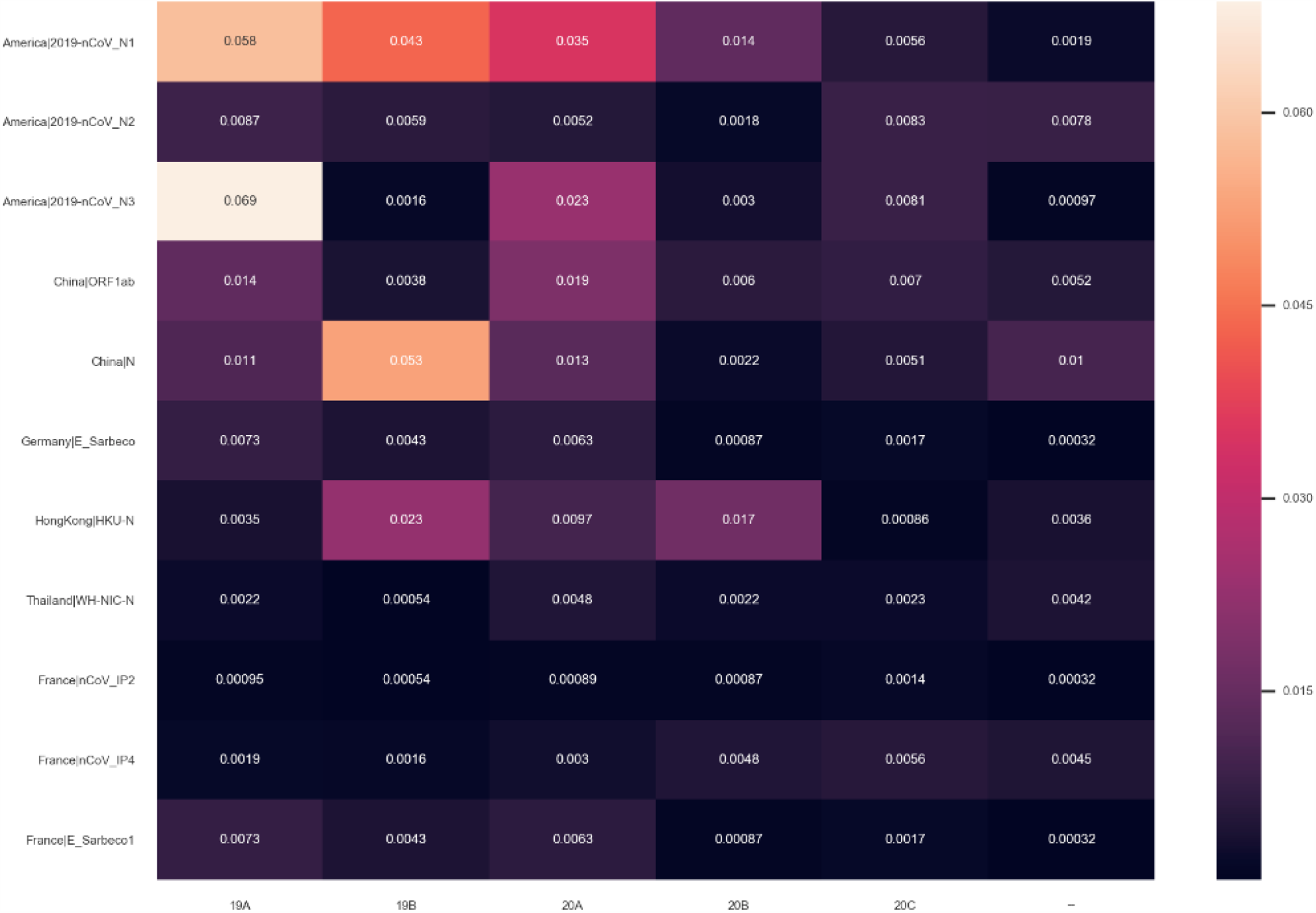
Average number of mutations for each PCR test that occur within each clade, as defined by NextStrain.

## Discussion

By taking a global perspective on both the SARS-CoV-2 genomes and the common RT-PCR protocols, we are able to highlight important trends within the data. We observe a an increasing number of mismatches between the primer and target genome sequence as time progresses. We can also see that the number of mismatches is higher when we compare genomes sampled outside of the country that designed the test compared to within the country. While these metrics do not quantify the performance of the test, they demonstrate a growing divergence between the targeted gene sequences and the test primers.

As shown by D. Bru et al.^12^, a single mutation can result in an underestimation of the gene copy number by up to 1000-fold. Our results reveal, today, an average of 1.1 mismatches between the primer and target sequences, with a growth of 2% each month. Understanding copy number is critical to correct interpretation of a PCR assay. If the genome being tested has sufficient mismatches this can lead to an erroneous copy number and, therefore, a misinterpretation of the assay result. In the case of SARS-CoV-2, for each targeted gene sequence, there are at least 10 different sequence variants and with this sequence diversity of the targeted genes, the mismatches in PCR primers may not be amplifying each example at the same rate, leading to false negatives. The given primers average a base length of 20 primers, and it has been demonstrated for primers with such base pair length that 2 to 3 mismatches reduces the yield by approximately 22 percent^13^. Our data indicates that this level of mismatches will be reached within 19 months or fewer if the rate of infection, and thus mutation, increases significantly.

The results of this study also demonstrate that each primer target develops a different number of mismatches over time (see: Figure 3). From the total number of mismatches created by primer target, we can see that the nucleoprotein targets from America, China, Hong Kong, and Thailand develop the greatest number of mismatches. Furthermore, when looking at the distribution of average number of mismatches over time, the primers targeting nucleoprotein have the largest distribution. The results indicate that primers targetting the envelope small membrane protein and the RNA-dependent RNA polymerase are the most resistant to mismatches. This may suggest more stable targets for future primer test designs.

The mutations that lead to mismatches between gene PCR primers and their targets reflect the sequence evolution of the virus. Comparing the difference in time of collection of two genomes with the number of mismatches by which they differ shows evidence for this evolution (Figure 4). Genomes that occur on the same day (delta time=0) have approximately zero difference, while genomes that occur at delta time=100 [days] have an average of 0.01 mismatches per nucleotide. This is consistent with the observed increasing number of mismatches over time, and shows that evolution of SARS-CoV-2 genomes is being sustained.

The continual branching of the genetic tree due to mutation is further supported by the analysis of the number of mutations within and outside the country that designed the particular primer. Figure 5 shows that most countries primers perform better when tested against genomes sequenced within the country rather than globally sequences genomes. In two cases, Hong Kong and France, the primers have a smaller percent of mismatches with genomes outside the country. For France, the IP2, a region of the RdRp gene, primer target creates a disproportionate number of mismatches when compared to genomes sequenced within France. This suggests that this region of the genome has deviated more from the original reference used to generate the primer set. For Hong Kong, they have the least number of genomes sequenced within the country in this dataset, so it is possible that the larger percent of mismatches for genomes within versus outside the country is an artifact of bias in data.

Nextstrain categorizes the various genetic phylogenies by clade, which is designed to denote long-term genetic changes based on mutation. Each clade defined requires significant geographical and frequency. This study shows that less than 3.5% of the regions on the genome that define the clades overlap with the region that the primers target. This indicates that variations in the primer target sequences have not yet have reached large enough statistical significance to define a new clade in the Nextstrain phylogeny, although the variants that are present in the primer region may cause a decrease in amplification signal within the assay.

With the emergence of specific mutations that are spreading at faster rates, this analysis becomes more important in evaluating the possible need for primer re-design. The emergence of the B.1.1.7 strain contains mutation in the regions encoding for the envelope small membrane protein and the nucleoprotein, both targeted by the current primers. With the number of cases of SARS-CoV-2 globally, it is highly probable that the genome will mutate in the primer target regions.

## Methods

### 5 Data Description

GISAID has emerged as a leading source of SARS-CoV-2 genomes, containing the largest number of genomes sequences around the world with metadata about the location and time of collection^14^. SARS-CoV-2 genomes from the GISAID repository were curated, collecting high quality genomes within the date range Aug 24, 2017 – July 31, 2020^15^. While this date range precedes the start of the current outbreak, the genome sequences from the earlier points and time serve as a control for comparison. We define high quality genomes as those with less than 1% N within the sequence and less 0.05% unique non-synonymous mutation. By taking these measures, we reduce the noise generated from random mutations or sequencing errors found within the genome. This resulted in a set of 61,996 SARS-CoV-2 genomes, for which we evaluated primer homology.

The WHO has published primers from six countries - China, France, USA, Japan, Germany, Hong Kong, and Thailand^5^. Each protocol published is a RT-PCR assay method, and for each primer set, a forward, reverse and probe sequence is provided^5^. For this study, we use the sequences as provided with no modifications made.

### 6 PCR Primer Comparison

Using the primer sequences and SARS-CoV-2 genomes described above, we perform a sequence comparison. Specifically, we used BLASTN with parameters similar to Primer-BLAST^16^. This procedure was verified to account for full alignments of the forward, reverse, and probe sequences of primers^17^. The BLAST results are then parsed, ensuring that the forward, reverse, and probe sequences match a given genome and that the probe sequence is matched spatially in the forward and reverse directions on the genome, and the number of mismatches is aggregated for each PCR sequence and genome. This metric does not necessarily predict whether the PCR test would generate a positive or negative outcome for the particular genome, but rather measures variability within the targeted gene region. Since all genomes included in this corpus are associated with SARS-CoV-2, its can be assumed that they were collected by a positive assay. Mutations in the targeted gene region, over time, can affect the sensitivity of the primers.

### 7 Time Analysis Methods

For each regional test, the primers each target a particular section of the genome derived from various reference genomes. However, as replication and mutation of the virus occurs, these targeted regions of circulating virus genomes accumulate sequence differences from the reference. Thus, the efficacy of the primer may decrease over time. As more mutations accumulate, it is important to measure the rate of mismatch growth between primer sequence and targeted section as a function of time. From this rate it is possible to anticipate when target sequences used in a regional test should be updated. To estimate the mutation rate of the targeted genes over time, we group the genomes by their date of sampling and aggregate the number of mismatches for each day. In order to reduce noise from days with few genomes collected, for any time-based analysis, we consider only those days that have over 100 unique genomes sequenced. With this restriction data is available for a time range between Jan 1, 2020 - July 25, 2020, for a total of 207 days. This process removes outlier data that was sequenced prior to the start of the pandemic, including sequences that were collected from non-human hosts.

### 8 Geographical Analysis Methods

As the virus has spread throughout the world, we see particular mutations that are specific to outbreaks by geospatial location. As studies using Bayesian coalescent analysis have shown, high evolutionary rates and fast population growth of the SARS-CoV-2 virus results in increasing diversification of the virus by geographic location^18^. To understand how the PCR tests respond differently for genomes collected by country, we first extract the country of sampling for each genome from the fasta header provided by GISAID and then group the number of mismatches found in the genome by in country versus out of country.

### 9 Clade Analysis Methods

SARS-CoV-2 genomes have been categorized into clades to define groups of mutations. For this analysis, we use the clades as indicated by NextStrain, which are defined by frequency and geographic spread. Their script to categorize genomes within the specific clade definitions was used to classify each genome within the dataset^19^. Furthermore, NextStrain publishes the genome locus that defines each clade, and these loci were compared to the genome location the primer targets bind to. By grouping the number of mismatches for each PCR by the genomes’ clade we see how different genetic variations affect the PCR test performance.

## Acknowledgements

The authors would like to acknowledge the GISAID Initiative and NCBI for the provision of data.

## Author contributions statement

G.N. conceived the experiment and analysis, M.K. verified the results, E.S. was the architect of the platform used, A.A and

K.L.B. performed genome quality analysis, J.H.K and V.M. provided scientific guidance and domain specific knowledge,

